# Performance evaluation of three DNA sample tracking tools in a whole exome sequencing workflow

**DOI:** 10.1101/2022.01.11.475818

**Authors:** Gertjan Wils, Céline Helsmoortel, Pieter-Jan Volders, Inge Vereecke, Mauro Milazzo, Jo Vandesompele, Frauke Coppieters, Kim De Leeneer, Steve Lefever

## Abstract

**Introduction:** Next-generation sequencing applications are becoming indispensable for clinical diagnostics. These experiments require numerous wet and dry lab steps, each one increasing the probability of a sample swap or contamination. Therefore, an identity confirmation at the end of the process is recommended to ensure the right data is used for each patient.

**Methods:** We tested three commercially available, SNP based sample tracking kits in a diagnostic workflow to evaluate their ease of use and performance. The coverage uniformity, on-target specificity, sample identification and genotyping performance were determined to assess the reliability and the cost-effectiveness of each kit.

*Results and discussion:* Hands-on time and manual steps are almost identical for the kits from pxlence and Nimagen. The Swift kit has an extra purification step, making it the longest and most demanding protocol. Furthermore, the Swift kit failed to correctly genotype 26 out of the 46 samples. The Nimagen kit identified all but one sample and the pxlence kit unambiguously identified all samples, making it the most reliable and robust kit of this evaluation. The Nimagen kit showed poor on-target mapping rates, resulting in deeper sequencing needs and higher sequencing costs compared to the other two kits. Our conclusion is that the Human Sample ID kit from pxlence is the most cost-effective of the three tested tools for DNA sample tracking and identification.

**Key points:**

- Kits from pxlence and Nimagen are easy to use.
- Unambiguous identification of all samples possible with the pxlence kit.
- Only 20 out of 46 samples were correctly identified with the Swift kit.
- Poor on-target rates for the Nimagen kit results in higher sequencing costs.

## 1. Introduction

Whole-exome (WES) and whole-genome sequencing (WGS) have become routine practice in clinical genetic laboratories [1]. However, the complex workflows, custody transfers and large datasets impose challenges on data integrity that range from the initial sample collection to the downstream data analysis. It is estimated that up to 3% of all samples may be compromised by provenance errors, raising serious concerns about the integrity and reliability of massively parallel sequencing (MPS) data [2–4]. Both in the clinic and the research laboratory, identity mix-ups can have detrimental consequences. A wrong diagnosis resulting in an incorrect or delayed treatment can cause severe harm to the patient, while erroneous data in a research context can impair discovery of new causal variants by yielding misleading variant candidates [5,6]. As sample mix-up errors are difficult to detect or to prevent, implementation of appropriate measures are critical for the unambiguous re-identification of samples throughout all stages of the MPS workflow [7,8]. An independent *post hoc* verification that the sequence results have been correctly assigned to each patient is therefore highly desirable.

Already in 2013, the American College of Medical Genetics and Genomics (ACMG) advised to track sample identity throughout the MPS process as part of adequate quality control [9]. The need for sample tracking was also included in more recent guidelines for (diagnostic) MPS issued by for instance the European Society of Human Genetics [7] and the Canadian College of Medical Geneticists (CCMG) [10]. Different methods exist for DNA sample tracking such as spike-in synthetic DNA standards [8,11,12] or single nucleotide polymorphisms (SNP) panels. By genotyping SNPs through an independent analysis, a unique fingerprint can be determined for each individual sample without interfering with the original DNA, ensuring sample mislabeling and handling errors are no longer part of the workflow [2,13–15]. Over the last years, several SNP-based sample identification panels specifically designed for MPS have been commercialized. In this study, a comparison of the performance of three commercially available SNP sample tracking methods is provided.

## 2. Materials and Methods

### 2.1 Patient samples

In total 46 different genomic DNA (gDNA) samples were used in this study, isolated from either blood (40 samples, MagCore Genomic DNA Large Volume Whole Blood Kit, MagCore Automated Nucleic Acid Extractor), Formalin-Fixed Paraffin-Embedded (FFPE) tissue (3 samples) and fresh frozen tissue (1 sample, QiaAmp Blood mini kit, QIAcube, QIAgen). Two samples are reference samples (NA24385 and NA12892) from the NIGMS Human Genetic Cell Repository at the Coriell Institute for Medical Research. For one FFPE donor, three biological gDNA replicates were included. DNA concentration and quality was determined with UV spectrophotometry (MagCore HF16 Super, RBC Bioscience). DNA concentration was higher than 30 ng/µl and the ratio of absorbance at 260 nm and 280 nm higher than 1.85 for each sample showing a succesful extraction and qualitative sample.

### 2.2. SNP sample tracking library preparation

The following commercially available SNP sample tracking kits were evaluated: Human Sample ID Kit PXL-SID-001 V1.0 (pxlence, kit A), Human Identification and Sample tracking kit RC-HEST V2.2 (Nimagen, kit B) and Accel-Amplicon Sample_ID Panel CP-UZ6128 V3.0 (Swift Biosciences custom panel, kit C, renamed as xGen Sample Identification Amplicon Panel after acquisition of Swift Biosciences by Integrated DNA Technologies). Since the execution of these experiments, pxlence has updated its kit. Characteristics of this version are equal or better than the previous version with reduced hands-on time due to the single-tube protocol (Figure B1). An overview of SNPs and gender markers (GM) present in each kit is given in Table A1, Table A2 and Table A3. All three kits were used as recommended by the manufacturer. A 20 ng/µl DNA dilution was made for each sample. Following the manual, different amount of input was used (kit A 20 ng, kit B 80 ng and kit C 20 ng). Quality control of the resulting library preparations was performed using concentration measurement (Fluoroskan, ThermoFisher, Invitrogen Quant-iT dsDNA Assay Kit, high sensitivity).

### 2.3. Sequencing

Per kit, library preps were pooled equivolumetrically, followed by bead purification (AMPure XP, Beckman Coulter) and concentration measurement of the final pools using qPCR (Kapa Library Quantification Kit, Roche). Subsequently, the three pools were spiked in a diagnostic WES workflow containing 186 exomes (SureSelectXT Low Input Target Enrichment System, Human All Exon V7 probes, Bravo Automated Liquid Handling Platform, Agilent), and 2 whole genome preparations (NEXTFLEX Rapid XP DNA-seq kit, PerkinElmer). The following ratios were applied for pooling: 1.27% for the sample tracking kits, 87.03% for the 186 exomes, and 11.70% for the 2 genomes. 1.19 nM of the final pool, including 1% PhiX, was sequenced on an Illumina NovaSeq 6000 system (S4 Reagent Kit, 300 cycles, paired-end sequencing).

### 2.4. Data Analysis

For assessing on-target specificity and coverage uniformity, reads were first aligned to the human reference genome (GRCh38) by means of the Burrows-Wheeler aligner (BWA v0.7.17) [16]. Mosdepth (v0.2.3) and total sample read-depth were used to calculate per-nucleotide normalized coverage to determine coverage uniformity of the various SNPs per patient [5]. To assess specificity, only regions having a non-normalized minimum per-nucleotide coverage of 25x and overlapping with a SNP included in the corresponding kit, were considered to be on-target. For analysing genotype similarities between WES and sample tracking data, individual libraries were downsampled to 100,000 reads. Genotype matches through logarithm of the odds (LOD) scores were used for comparison of genetic fingerprints between samples using the CrosscheckFingerprints tool from the Picard software package (v2.1.1) [17]. In this analysis, a near zero LOD score indicates an inconclusive comparison, while a sample match or mismatch are given a positive or negative LOD score, respectively. LOD values greater than 5 were considered a match, lower than -5 as a mismatch. Values between -5 and 5 were labeled as inconclusive. Data was filtered and matrices made with R (v4.1.2). Gender determination differed for each kit. For kit A, genetic fingerprints were made for the SNP located on the Y chromosome as described above. Normalised coverage for Amelogenin X and Y was used as an additional control. The normalised coverage on Amelogenin X and Y was used to determine gender for kit B. For kit C, we looked at the median normalised SNP coverage compared to the normalised coverage on chromosome X.

## 3. Results and discussion

### 3.1. Wet-lab procedure and sequencing output

Sample preparation, clean-up and quantification is very similar for the tested kits. The samples were processed in parallel using the different kits (Figure 1). Kit B has the shortest hands-on time (30 min) but required four times the amount of DNA input (80 ng) compared to the other kits. The PCR is a one-tube reaction which makes the sample preparation straightforward and easy. The hands-on time for kit C is roughly 1 hour and 30 minutes. This is because the indexed sequencing adapters are added separately after the multiplex PCR. In between, magnetic beads (AMPure XP, Beckman Coulter) are used for a size selection clean-up. These extra steps in the protocol allow for a lower input (10-25 ng) but increase the hands-on time drastically. Kit A has a hand-on time of 1 hour. The reaction is split into two steps similar to kit C. A part of the SNP-enriched DNA is added to the indexing assay which results in the final product ready for clean-up and sequencing. Of note, a single-tube version of kit A is recently made available, and decreases the hands-on time to less than 30 minutes. Characteristics, like sensitivity and uniformity, of this improved protocol are equal to the previous version (supplementary Figure B1). DNA input for kit A can range from 2 to 20 ng, making it the most sensitive kit in this evaluation. The average library concentration before clean-up was 150.2 nM for kit A, 1063.5 nM for kit B and 142.2 nM for kit C. There was no significant difference between PCR yield of blood, FFPE or fresh frozen tissue samples (p>0.05) showing that each kit can be used with all three types of samples although further testing should confirm this as only three FFPE and one fresh frozen tissue sample were included in this study. Concentration of the sample pools after clean-up was 406.4 nM for kit A, 1109.3 nM for kit B and 49.8 nM for kit C. Sequencing yield significantly varied: kit A resulted in 2.4xE19 reads/nmol, 3.0xE18 reads/nmol for kit B and 4.9xE18 reads/nmol for kit C (p<1E-4). This shows concentration determination is most accurate with kit A. Median SNP coverage per 2000 reads per DNA sample was calculated to be 23 for kit A, 22 for kit B and 35 for kit C. Kit C has the greatest SNP coverage, due to the size of the panel that only contains 28 SNPs. Median SNP coverage per 2000 reads for blood samples and FFPE samples was not significantly different for kit A and B. FFPE samples with kit C resulted in a lower number of reads compared to the blood samples (p=0.0028) indicating that amplification did not result in usable amplicons or clustering of the SNP panel was less efficient for these lower quality samples.

**Figure 1.**
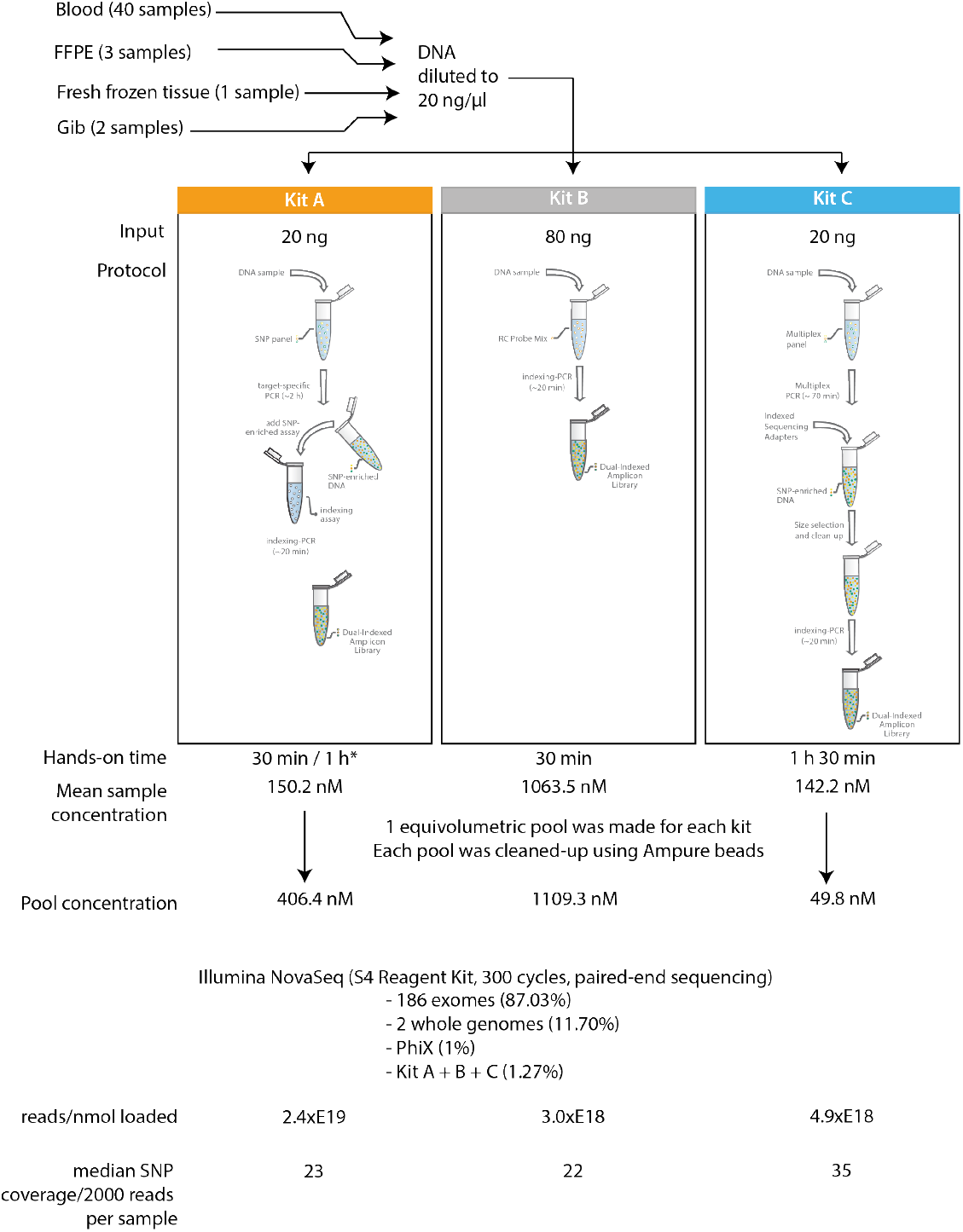
Overview of experimental setup and protocols. Data on hands-on time, mean sample concentration after PCR, concentration of the sample pool after clean-up, number of reads per nmol pool loaded, median SNP coverage per 200,000 reads per sample is shown. * version 1.0 of kit A was used for this evaluation. Version 1.2 has a workflow similar to kit B with a hands-on time of 30 min.

### 3.2. Gender determination

Kit A includes six gender markers, five of them are located on the Y chromosome and one on AMELX/Y. This primer pair results in an amplification on both the X and Y chromosome with a difference in amplicon sequence and length. This enables robust discriminate between genders even with lower quality or degraded DNA. Kit B contains two primer pairs amplifying the amelogenin gene on either the X and Y chromosome. Kit C only contains one SNP located on the X chromosome, making it the least reliable option for gender determination. Gender was correctly determined by kit A in all samples. Kit B determined 43 samples correctly, 2 were marked as inconclusive and 1 was assigned the wrong gender. Kit C had 6 inconclusive samples and 1 miss-identified gender. This clearly shows that a sufficient number of markers is required for accurate determination of gender.

### 3.3. Coverage uniformity

Sequencing coverage uniformity is a measure of the amplification efficiency of each individual SNP within the multiplex PCR reactions performed as part of each of the corresponding method’s workflow. Perfect equimolar assay coverage means minimal sequencing capacity is required to attain a minimal coverage per assay (for example, in this setting each assay would be covered exactly 30 times), resulting in an optimal cost-efficiency. Deviation of such a perfect situation results in increased sequencing capacity – and sequencing cost – required to achieve similar results (e.g. if a method includes a sub-performing assay, additional sequencing capacity would be needed to bring that assay up to 30x coverage). Coverage uniformity is typically reported as the percentage of assays having a coverage above 0.2 times the median coverage in a specific sample. However, since this measure ignores highly-efficient assays with excessive coverage – resulting in decreased coverage uniformity – the percentage of assay falling within the range of 2-fold around the median sample coverage (calculated across all samples) is also determined here. Gender markers were omitted from this analysis because of unequal coverage between male and female samples. Results indicate that kit B scores best on coverage uniformity, with 90.58% of the datapoints within 2-fold of the median and is significantly different from the two other kits (p < 1E-04) (Figure 2). It is closely followed by kit A and kit C, with 83.52% and 81.24% of the datapoints within a 2-fold range around the median, respectively (Figure 2). These findings are confirmed when calculating the per-sample standard deviation (SD) of the normalized coverage: 0.0029 for kit B, 0.0040 for kit A, and 0.0076 for kit C (Figure 3A). Remarkably, kit B shows a much larger inter-sample intra-assay variability compared to the other two methods tested (as reflected by the wide box plots).

**Figure 2.**
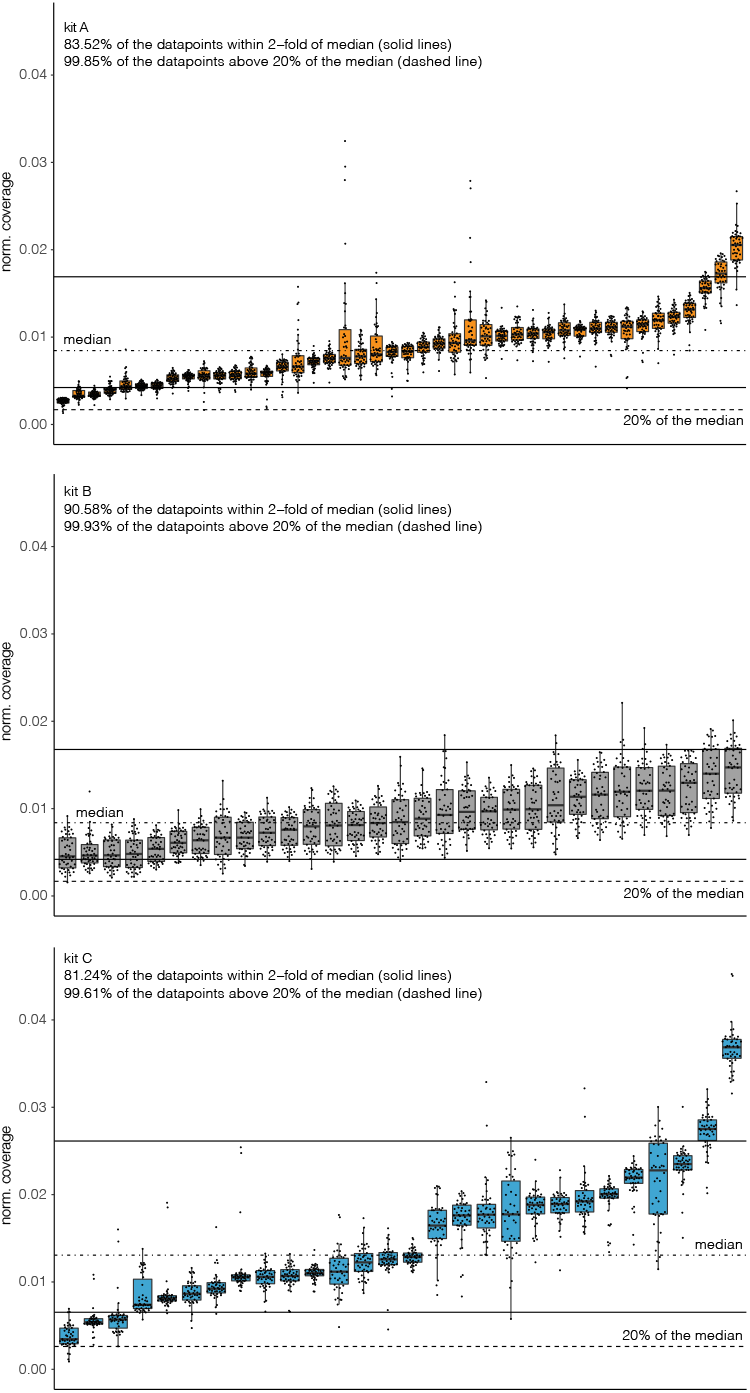
Normalized sequence coverage on the regions of interest across all samples for each of the kits. Dotted-dashed line indicates the median normalized coverage across all datapoints, solid lines indicate the upper- and lower threshold of the 2-fold of the median range, and dotted line indicates the 20% of the median coverage (each dot is a patient; each boxplot is a SNP).

**Figure 3.**
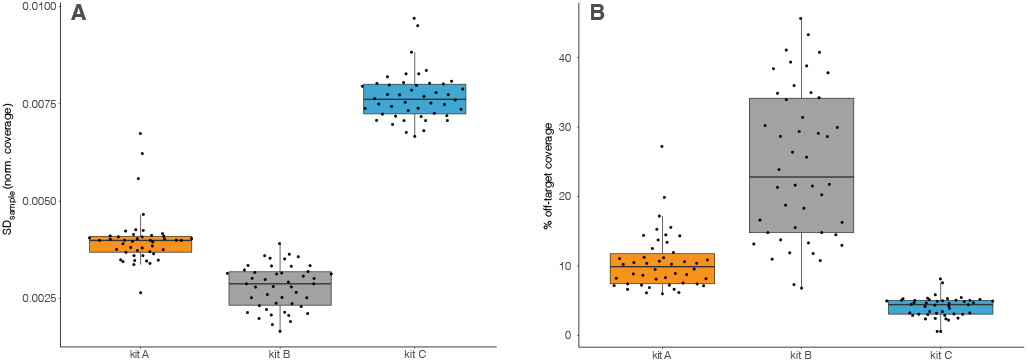
(A) Standard deviation of the normalized coverage per sample across all regions of interest for each of the kits, (B) percentage of off-target coverage per sample for each of the kits (lower is better; each dot is a patient).

### 3.4. On-target specificity

Analogous to coverage uniformity, on-target specificity has an impact on the sequencing capacity required per sample, and thus per-sample sequencing cost. In an ideal scenario, all reads produced by a method will generate useful data for the targets of interest. A lower on-target specificity – and thus higher level of non-specificity – will lead to lower coverage for the SNPs thus a need for higher per-sample sequencing capacity to achieve the same minimal per-assay coverage. This analysis shows that kit C performs best in the context of on-target specificity, with a median of 4.43% of the reads aligning off-target (Figure 3B). Kit A also scores very well with approximately 9.90% off-target reads. Kit B performs worst with a median of 22.82% and a significant portion of the samples showing off-target percentages above 30% (n = 11) and even up to 40% (n = 4). A much smaller range in per-sample off-target percentages was observed for both kit A and kit C. This means that, in comparison to kit A and kit C, kit B will need up to 30-35% more per-sample sequencing capacity for a significant portion of the samples, resulting in an overall higher per-sample sequencing cost.

### 3.5. Genotyping and sample discrimination

LOD scores were used for comparing the genotypes obtained by the three SNP sample tracking panels and the data from the WES (Figure 4A). Only for kit A, unambiguous sample discrimination and identification was obtained for all samples. For kit B, one genotyping sample was not identified correctly, showing as a mismatch with the correct WES data, and being inconclusive with another unrelated sample. With kit C, only 32 of the 46 samples were matched with its corresponding WES data and -more importantly -only 20 samples showed a correct match while not having an inconclusive result with another sample. For none of the three kits a match was found with an unrelated sample, indicating that no sample mix-ups occurred. The mean fraction of correctly called SNPs per sample is 99.2 % for kit A, 99.8 % for kit B and 99.4 % for kit C. The excellent performance of kit A – and to a lesser extent kit B – is further substantiated by looking at the individual LOD scores of the matching samples and mismatch samples (Figure 4B). For a good discriminatory performance, LOD scores should be as decisive as possible, meaning LOD scores of matching samples should be as high as possible above zero and LOD scores of mismatch samples should be as low as possible below zero. As expected from the genotyping, kit C shows the lowest discriminatory performance, with average LOD scores of ±6 being just above the inconclusive threshold. In comparison, LOD scores of kit A and kit B showed a much better discrimination with LOD scores of the matching samples of respectively ∼19 and ∼13, and LOD scores of mismatch samples of ∼50 and ∼-40, respectively. The overall excellent performance of kit A in discriminating samples can be in large part explained by the larger number of correctly called SNPs in this kit. A larger number of assays has the additional benefit that – in case of sub-optimal sequencing with lower coverage values – sufficient high-quality markers remain for robust sample identification. In contrast, kits with lower SNP numbers can suffer from lack in discrimination power when combined with sub-optimal sequencing results due to insufficient high-quality markers remaining.

**Figure 4.**
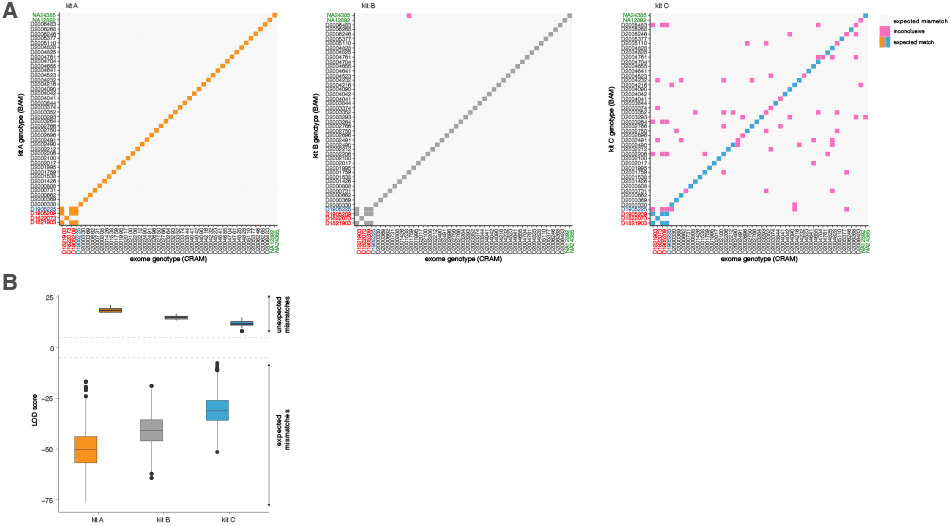
(A) Match and mismatched samples between the exome data (X-axis) and the SNP genotypes obtained with the sample ID kits (Y-axis), (B) LOD scores for the expected matches and unexpected mismatches for each of the kits (more extreme values are better) FFPE sample names are coloured red, the fresh frozen tissue sample blue and reference samples in green. Sample D1821903, D1822073 and D1905225 are biological gDNA replicates.

## 4. Conclusions

In a real-life clinical setting, the three tested SNP sample tracking methods displayed significant differences in their sample identification and genotyping performance (Table 1). Overall, kit C showed to be unreliable with many samples that showed undecisive correlations, although on-target specificity was observed to be highest in this kit. Kit B performed best on coverage uniformity but showed poor on-target rates resulting in higher sequencing costs. From the three kits, kit A excelled in sample identification and discrimination. The high sample discrimination performance allows for a high-confident and robust genotyping, assuring correct sample identification and avoids the need for reanalysing samples. Combined with an above-average on-target specificity and coverage uniformity, kit A shows the overall best per-sample cost-efficiency. When taking into account the version 1.2 of kit A, hands-on time and number of manual steps is identical between kit A and kit B. Kit C has a second PCR step and additional clean-up step, making it a more time intensive protocol. As this evaluation was the start of the implementation of a sample tracking solution for the Center for Medical Genetics in Ghent, no reference protocol was available at the time to compare the above-mentioned findings. While this evaluation included some FFPE and fresh frozen tissue samples, further testing should be performed to confirm the effectiveness for these sample types.

**Table 1.** Overview of kit characteristics and performance parameters evaluated in this study (best score in bold)

|  | Human Sample ID kit (pxlence, kit A) | Human Identification and Sample tracking kit (Nimagen, kit B) | Accel-Amplicon Sample ID Panel (Swift Biosciences, kit C) |
| --- | --- | --- | --- |
| DNA input range (ng) | **2 – 20** | 80 – 100 | 10 – 25 |
| number of SNPs | **44** | 34 | 28 |
| number of gender markers | **6** | 2 | 1 |
| number of reads / nmol loaded | **2.4xE19** | 3.0xE18 | 4.9xE18 |
| median SNP coverage / 2000 reads / sample | 23 | 22 | **35** |
| coverage > 20% of mean | 99.85% | **99.93%** | 99.61% |
| coverage within 2-fold of median | 83.52% | **90.58%** | 81.24% |
| standard deviation of coverage | 0.0040 | **0.0029** | 0.0076 |
| median % off-target | 9.90% | 22.82% | **4.43%** |
| median LOD match (more is better) | **19** | 13 | 6 |
| median LOD mismatch (less is better) | **-50** | -40 | -31 |
| conclusive genotypes | **46/46 (100%)** | 45/46 (98%) | 20/46 (43%) |

## Statements and declarations

### Author Contributions: Conceptualization

Jo Vandesompele, Steve Lefever and Frauke Coppieters; methodology, Inge Vereecke and Mauro Milazzo; software, Pieter-Jan Volders and Steve Lefever; validation, Inge Vereecke and Mauro Milazzo; formal analysis, Pieter-Jan Volders, Steve Lefever and Gertjan Wils; investigation, Mauro Milazzo and Inge Vereecke; writing—original draft preparation, Gertjan Wils; writing—review and editing, Frauke Coppieters, Jo Vandesompele, Mauro Milazzo, Steve Lefever, Kim De Leeneer and Gertjan Wils; visualization, Pieter-Jan Volders, Steve Lefever and Gertjan Wils; supervision, Jo Vandesompele; project administration, Céline Helsmoortel; All authors have read and agreed to the published version of the manuscript.

### Informed Consent Statement

Patient consent was waived due to the experiments not adversely affecting the rights and welfare of the subjects.

### Data Availability Statement

The data that support the findings of this study are available on request from the corresponding author. The data are not publicly available because they contain information that could compromise research participant privacy/consent.

