## Supplementary material for "Performance evaluation of three DNA sample tracking tools in a whole exome sequencing workflow"

### Appendix A

**Table A1** Overview of SNPs and gender makers (GM) in the Human Sample ID Kit PXL-SID-001 V1.0 (pxlence, kit A). Additional information can be found on https://www.pxlence.com/products_sample_tracking.php.

| SNP and GM nr | reference SNP ID | hg38-coordinates | |
|---|---|---|---|
| | | chr | position |
| SNP1 | rs1410592 | 1 | 179551371 |
| SNP2 | rs2076356 | 1 | 209638541 |
| SNP3 | rs2013162 | 1 | 209795339 |
| SNP4 | rs2229267 | 2 | 169235885 |
| SNP5 | rs1560221 | 2 | 178589667 |
| SNP6 | rs2163009 | 2 | 178590480 |
| SNP7 | rs10498027 | 2 | 214955289 |
| SNP8 | rs3738985 | 2 | 44275649 |
| SNP9 | rs4688963 | 4 | 5748177 |
| SNP10 | rs2736982 | 4 | 87613083 |
| SNP11 | rs4669 | 5 | 136056737 |
| SNP12 | rs3088052 | 5 | 139121126 |
| SNP13 | rs7823 | 5 | 54456158 |
| SNP14 | rs10265207 | 7 | 33970334 |
| SNP15 | rs7738 | 7 | 43807004 |
| SNP16 | rs3808554 | 8 | 103324868 |
| SNP17 | rs3124768 | 9 | 133439376 |
| SNP18 | rs639225 | 9 | 27202872 |
| SNP19 | rs7859201 | 9 | 74800368 |
| SNP20 | rs6163 | 10 | 102837167 |
| SNP21 | rs2673794 | 10 | 68166340 |
| SNP22 | rs1131824 | 10 | 77184832 |
| SNP23 | rs17109674 | 10 | 94032006 |
| SNP24 | rs1043388 | 11 | 6608435 |
| SNP25 | rs60637 | 12 | 51806958 |
| SNP26 | rs7300444 | 12 | 884764 |
| SNP27 | rs3742165 | 13 | 24892817 |
| SNP28 | rs9532292 | 13 | 38859469 |
| SNP29 | rs7161192 | 14 | 64170429 |
| SNP30 | rs2296409 | 16 | 68679827 |
| SNP31 | rs2296408 | 16 | 68679920 |
| SNP32 | rs17715450 | 16 | 68695882 |
| SNP33 | rs3762171 | 16 | 70512331 |
| SNP34 | rs2285479 | 17 | 10632701 |
| SNP35 | rs2285475 | 17 | 10639154 |
| SNP36 | rs5910 | 17 | 44372421 |
| SNP37 | rs1052706 | 17 | 73196524 |
| SNP38 | rs9962023 | 18 | 23833905 |
| SNP39 | rs2298628 | 18 | 49929553 |
| SNP40 | rs2228611 | 19 | 10156401 |
| SNP41 | rs11084673 | 19 | 32862558 |
| SNP42 | rs10373 | 20 | 6119441 |
| SNP43 | rs2249057 | 21 | 46353189 |
| SNP44 | rs4820268 | 22 | 37073551 |
| GM1 | AMELX/Y | X/Y | |
| GM2 | KDM5D | Y | |
| GM3 | SRY | Y | |
| GM4 | TXLNGY | Y | |
| GM5 | USP9Y | Y | |
| GM6 | UTY | Y | |

**Table A2** Overview of SNPs and gender makers (GM) in the Human Identification and Sample tracking kit RC-HEST V2.2 (Nimagen, kit B). Additional information about the kit can be found at https://www.nimagen.com/shop/products/rc-hest096/human-exome-sample-tracking-and-identification-kit-96-rxn.

| SNP and GM nr | reference SNP ID | hg38-coordinates | |
|---|---|---|---|
| | | chr | position |
| SNP1 | s1410592 | 1 | 179551371 |
| SNP2 | rs2229546 | 1 | 67395837 |
| SNP3 | rs10203363 | 2 | 227032260 |
| SNP4 | rs2819561 | 3 | 4362083 |
| SNP5 | rs4688963 | 4 | 5748177 |
| SNP6 | rs309557 | 5 | 83538811 |
| SNP7 | rs4735258 | 8 | 93923709 |
| SNP8 | rs4870723 | 8 | 120216440 |
| SNP9 | rs7465584 | 8 | 123975238 |
| SNP10 | rs1381532 | 9 | 97428498 |
| SNP11 | rs1536928 | 9 | 122629130 |
| SNP12 | rs1572983 | 9 | 101371346 |
| SNP13 | rs577993 | 9 | 76706955 |
| SNP14 | rs10883099 | 10 | 98459557 |
| SNP15 | rs4617548 | 11 | 16111867 |
| SNP16 | rs7300444 | 12 | 884764 |
| SNP17 | rs495680 | 13 | 33129519 |
| SNP18 | rs9532292 | 13 | 38859469 |
| SNP19 | rs11158685 | 14 | 67575857 |
| SNP20 | rs4577050 | 15 | 34236747 |
| SNP21 | rs1026128 | 17 | 73200670 |
| SNP22 | rs1037256 | 17 | 73201609 |
| SNP23 | rs1292053 | 17 | 59886176 |
| SNP24 | rs2159132 | 17 | 14102122 |
| SNP25 | rs1805034 | 18 | 62360008 |
| SNP26 | rs3826616 | 18 | 63987229 |
| SNP27 | rs9962023 | 18 | 23833905 |
| SNP28 | rs10373 | 20 | 6119441 |
| SNP29 | rs2296241 | 20 | 54169680 |
| SNP30 | rs4148973 | 21 | 42903480 |
| SNP31 | rs760482 | 22 | 38782696 |
| SNP32 | rs2073787 | X | 110451457 |
| SNP33 | rs5930933 | X | 136349199 |
| SNP34 | rs6568050 | X | 112454808 |
| GM4 | AMELX | X | |
| GM5 | AMELY | Y | |

**Table A3** Overview of SNPs and gender makers (GM) in the Accel-Amplicon Sample_ID Panel CP-UZ6128 V3.0 (Swift Biosciences custom panel, kit C). Additional information about the kit can be found at https://www.eu.idtdna.com/pages/products/next-generation-sequencing/workflow/xgen-ngs-amplicon-sequencing/predesigned-amplicon-panels/sample-id-amp-panel.

| SNP and GM nr | reference SNP ID | hg38-coordinates | |
|---|---|---|---|
| | | chr | position |
| SNP1 | rs1130683 | 1 | 11,072,160 |
| SNP2 | rs14291 | 2 | 36,583,443 |
| SNP3 | rs1004814 | 2 | 217,847,583 |
| SNP4 | rs2340917 | 3 | 14,133,762 |
| SNP5 | rs17610219 | 4 | 146,903,712 |
| SNP6 | rs2304035 | 5 | 168,749,512 |
| SNP7 | rs1126476 | 6 | 39,080,715 |
| SNP8 | rs2074603 | 7 | 11,541,507 |
| SNP9 | rs3735801 | 8 | 140,451,167 |
| SNP10 | rs2787374 | 9 | 100,292,669 |
| SNP11 | rs1043836 | 9 | 113,373,918 |
| SNP12 | rs1801222 | 10 | 17,114,152 |
| SNP13 | rs17109674 | 10 | 94,032,006 |
| SNP14 | rs4453265 | 11 | 74,074,281 |
| SNP15 | rs879732 | 12 | 12,087,265 |
| SNP16 | rs1061472 | 13 | 51,950,352 |
| SNP17 | rs2069540 | 14 | 23,433,544 |
| SNP18 | rs2470911 | 15 | 44,754,936 |
| SNP19 | rs2272554 | 16 | 15,756,347 |
| SNP20 | rs2296405 | 16 | 68,687,567 |
| SNP21 | rs9898751 | 17 | 8,047,634 |
| SNP22 | rs3737353 | 18 | 5,956,239 |
| SNP23 | rs3810198 | 19 | 16,490,383 |
| SNP24 | rs2071327 | 19 | 35,831,368 |
| SNP25 | rs611847 | 20 | 3,703,375 |
| SNP26 | rs6093935 | 20 | 44,114,814 |
| SNP27 | rs7278737 | 21 | 14,109,044 |
| SNP28 | rs4822360 | 22 | 23,140,273 |
| GM1 | rs3213466 | X | 153,345,527 |

### Appendix B

Experimental setup: 10 DNA samples (extracted from human blood) were used to compare version 1.0 and 1.2 of the Human Sample ID kit from pxlence in a different independent experiment. PCR was performed as described in the manuals. In short v1.0 consists of two PCR steps (SNP amplification and indexing) and v1.2 combines these two steps into a single step. Library preps were equivolumetrically pooled before bead purification (AMPure XP, Beckman Coulter) and concentration measurement of the final pools (one for each kit version) using qPCR (Kapa Library Quantification Kit, Roche). Pools were sequenced with 10% PhiX on an Illumina Miseq (Miseq Nano reagents kit v2, 300-cycles, paired-end sequencing).

Results and conclusion: Reads were equally distributed between the two versions showing that cluster efficiency is similar (Figure B1 panel A). Percentage on-target reads is slightly higher for version 1.0, 96% on average, compared to v1.2 (92%). This difference, although significant is rather small and will not result in a notable difference in sequencing cost (Figure B1 panel B). The percentage of SNPs withing 2-fold of the median coverage is 79% for v 1.0, this is inline with our findings during the evaluation of the three sample tracking panels considering the differences in samples and sequencing technology (Figure 2). The new version has a higher percentage of SNPs within 2-fold of the median (88%), showing an improvement on the previous version (Figure B1 panel C). SNP coverage is equimolar for both versions as shown in figure B1 panel D. The change to a one step, closed-tube protocol cuts the hand-on time in half, making v1.2 more user friendly with the same excellent performance (figure B1 panel E).

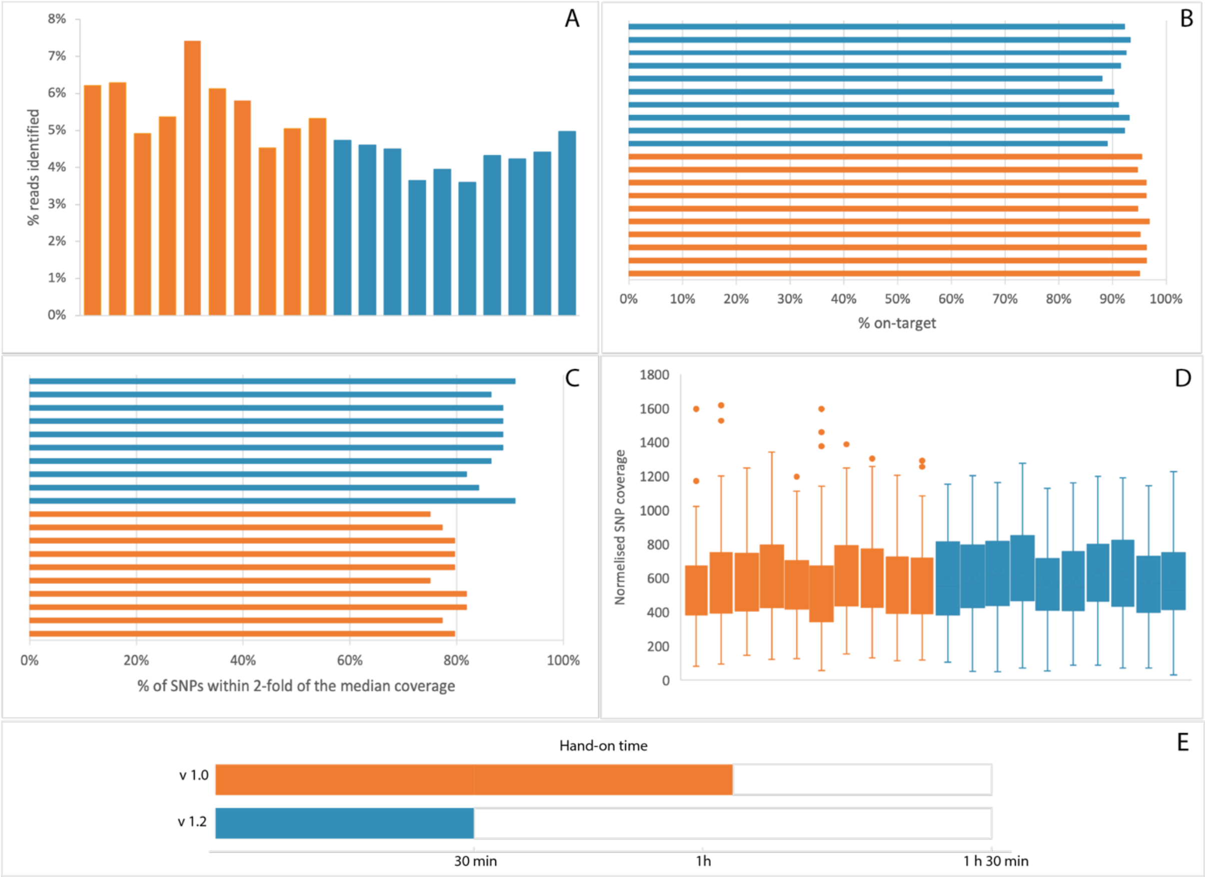


**Figure B1** Comparison between performance characteristics of Human Sample ID kit version 1.0 (orange) and v1.2 (blue) from pxlence. 10 samples were used in this experiment. Panel A: percentage of reads identified, Panel B; percentage on-target read, Panel C: percentage of SNPs within 2-fold of the median coverage, Panel D: Normalised SNP coverage
